# Live-HPF CLEM using the HPM Live µ: Finding back all needles in every haystack. The last frontier in CLEM

**DOI:** 10.1101/2024.10.21.619370

**Authors:** Xavier Heiligenstein, Chie Kodera, Yann Bret, Frederic Eyraud, Vincent Muczynski, Jérôme Heiligenstein, Martin Belle

## Abstract

Correlative light and electron microscopy is a scientific method that encompasses several technologies and workflows. One of the pioneering workflows consisted in vitrifying by high pressure freezing a sample, shortly after live observation^1^. Despite its extraordinary potential, it did not turn into a routine technique for practical reasons. We redesigned the entire tool set, from the sample carrier to the high-pressure freezing machine, to standardize and democratise the technique. In our manuscript, we present all the technological developments that lead to a routine workflow for live to HPF CLEM. We demonstrate our ability to track rapidly moving endosomes live and retrieve them at the electron microscopy level, in three dimensions, with high confidence.

## Introduction

Over the past two decades, correlative light and electron microscopies (CLEM), have become crucial for deciphering biological complexities across scales. A pivotal aspect of this workflow is the immobilization of samples in their near-native state with minimal artifacts, ensuring biologically relevant interpretations. Electron microscopy, with its extensive expertise and unparalleled resolution, is now challenged to transition from chemical fixation to physical cryo-immobilization via vitrification.

From the late seventies, various vitrification methods surfaced, but the two predominant techniques to date are “plunge freezing,” uniquely suited for isolated particles, and “high pressure freezing” (HPF ^2–7^), designed for samples ranging from individual cells to tissue sections as thick as 200 µm. HPF, in particular, has historically been perceived as intricate and unreliable. The rapid release of liquid nitrogen at a staggering 2000 bars, impacting the sample in less than 25 milliseconds, with a sustaining duration over 150 milliseconds, resembles an explosion that could easily disperse unprotected biological material. However, serving as a gateway to elaborate electron microscopy protocols (such as CEMOVIS^8–10^, Freeze-Substitution^11–15^, cryo-EM FIB-milling^16–18^), HPF has now become an indispensable technique for preparing samples with optimal ultrastructure preservation including multiscale tissues.

To date, most CLEM approaches relying on HPF, start by the cryo-immobilization, followed by light microscopy on either the vitreous sample ^17,19–21^ or the resin embedded sample (In Resin Fluorescence ^15,22,23^). Both workflows therefore lack the dynamic information.

Employing live imaging prior to high pressure freezing is an attractive idea to capture living dynamics in close to native state. However, it imposes a specific preparation, appropriate to optimise both steps. The biological specimen needs to be located within a device that facilitates live imaging in physiological conditions, while offering physical shielding against the high-pressure flow of liquid nitrogen. Furthermore, the temporal gap between these two steps must be minimized to remain relevant to physiological timescales ^1,24^.

To tackle this challenge, we pioneered the development of the CryoCapsule in 2013^25,26^. This innovative container comprises two sapphire discs separated by a spacer ring, offering ample volume for cultivating cell monolayers or deposing small model organisms or tissue biopsies. Its integrated structure optimizes the transition time from a light microscope to a High Pressure Freezer. Nonetheless, this transfer process remained a manual endeavour, contingent upon the operator’s skill to achieve reproducible outcomes, with the best performance clocking in at 15 seconds. To mitigate the operator’s influence, we introduced a groundbreaking amalgamation of light microscope and high-pressure freezer, resulting in the HPM Live µ^24^. This fusion of the HPM Live µ, the CryoCapsule and a light microscope has now reduced the reliable transfer time to 1.26 seconds. Precise time measurements span from the operator’s directives for vitrification initiation to the commencement of the vitrification process. This confluence of automation and advanced CryoCapsule technology has exponentially elevated the success rate of sample retrieval, nearly guaranteeing a 100% success rate. As a result, Live-CLEM-HPF has transitioned from an aspiration to a routine reality.

## Material and Method

### Setup

The HPM Live µ marks the latest advancement in the lineage of Baltec’s historical HPM010. It incorporates patented pressurizing technologies (patent filed PCT/EP2019071787) and an innovative graphical user interface (GUI) ^24^. Designed for routine HPF experiments on traditional carriers (3 and 6mm, membrane carriers, hats), tubes and the CryoCapsule, and offers seamless integration with a light microscope. This integrated setup positions the microscope beneath a robotic arm responsible for transferring samples into the HPF chamber. (Figure 1, Automatic transfer arm)

**Figure 1:**
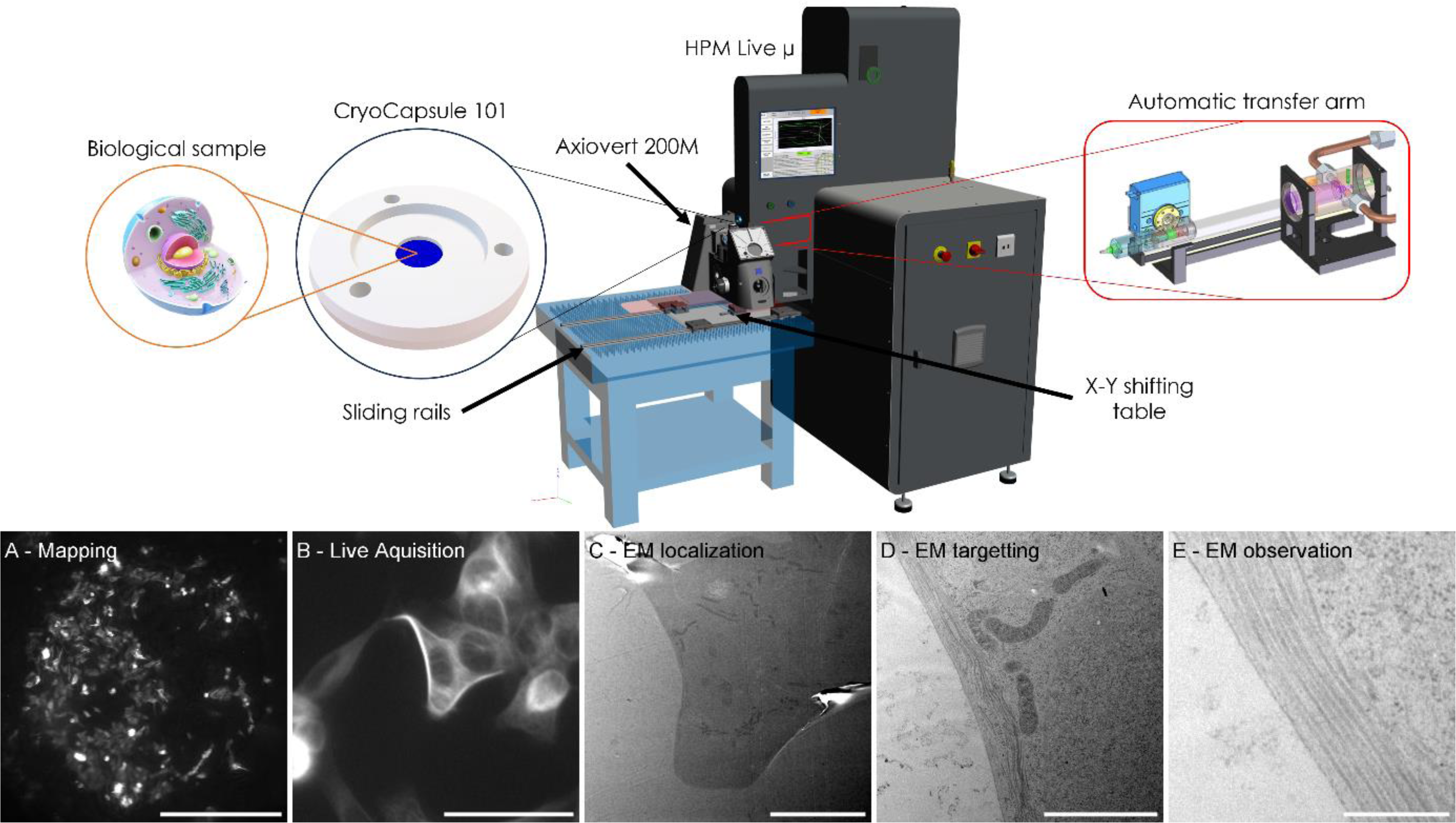
Workflow. The sample is cultured in a CryoCapsule type 101, and loaded at the tip of the robotic arm right above the microscope lens. Upon triggering, the sample is transferred in the HPF chamber for vitrification in 1.26 seconds. A-The sample is mapped live in the CryoCapsule. B-live imaging at 50x. C-Localization in TEM at low magnification. D-Targetting of the structure of interest. E-Observation of the structure of interest.

Through this configuration, we observe specimens continuously, granting visibility up to 1.26 seconds prior to vitrification. It ultimately offers to seamlessly image a sample from 5x mapping and 50x live observation (Figure 1 A, B, Figure 2 D, E, H, I) to electron microscopy localization, targeting and observation of the structure of interest (Figure 1 C-E). In the work presented here, we integrated a Zeiss Axiovert 200M with our HPM Live µ (Figure 1, Figure 2). In other laboratories, we successfully integrated the HPM Live µ with a Zeiss LSM900 Airyscan and a Nikon TiEclipse and Te2000. Operating on an X-Y shifting table, the microscope offers the ability to focus sharply on specific areas of live samples at higher magnifications (video 1). This design emulates the fixed-stage microscopes often employed in electro-physiological experiments.

**Figure 2:**
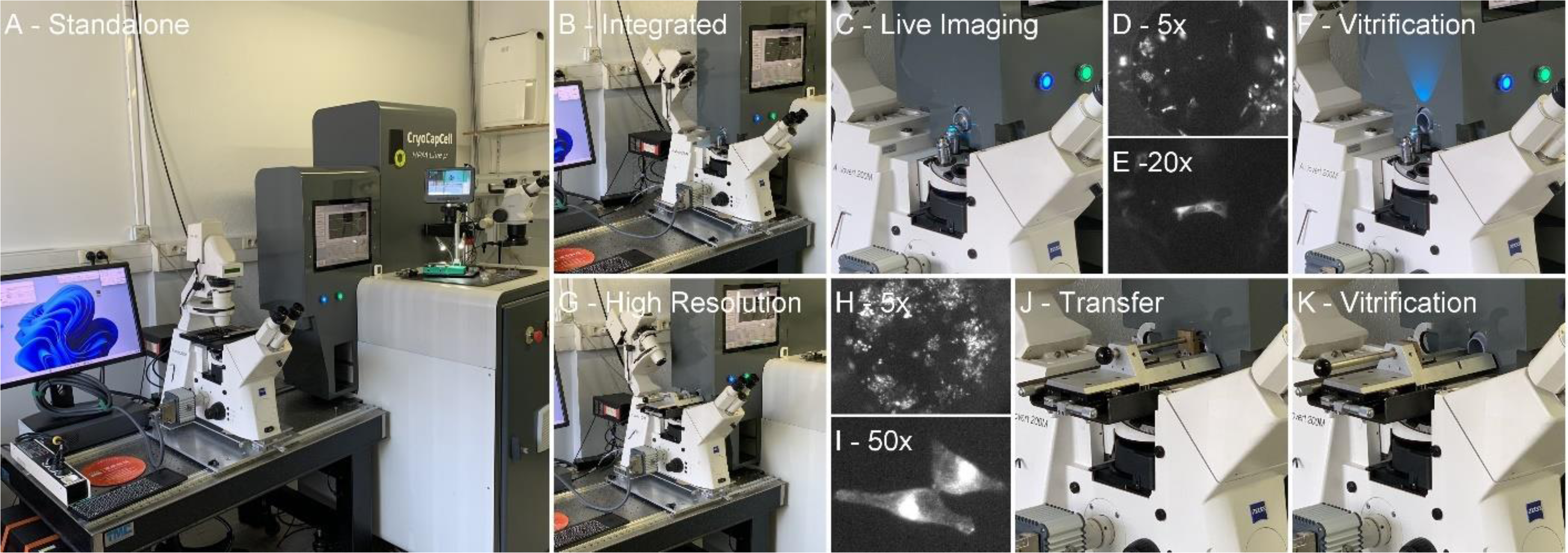
various microscope/HPF configurations. A the microscope is separated from the HPM for a standalone use. The HPM can also be used without the microscope. B-Integration of the microscope to the HPM. C-Live imaging. D-Low magnification observation at 5x to locate the cells of interest. E-Imaging at 20x. F-Automatic transfer for vitrification in 1.26 seconds. G-Integration of the microscope with the high-resolution table. H-Mapping at low magnification and centering of the cells of interest in the field of view. I-High magnification live imaging at 50x. J-Manual push of the sample holder into the HPM live µ. K-Triggering of the HPF after transfer (J) to initiate the vitrification in 1.26 seconds.

To enhance operational efficiency, both the X-Y shifting table and the microscope are positioned on sliding rails, allowing their separation from the HPM for individual use as stand-alone microscopes or HPM only (Figure 2-A). This adaptability ensures that users can optimize the use time of their microscope or HPM. The transition between the two modes—stand-alone and integrated—occurs swiftly, taking about a minute to complete (Video 1).

The microscope used in this study is a Zeiss Axiovert 200M, equipped with a Lumencor Light Engine AURA light source, an ORCA ER Hamamatsu camera. The whole apparatus is controlled by Micromanager 2.0.1 64bits^27–29^. List of objective lenses: 5x Fluar 0.25NA, 10x EC plan NeoFluar 0.3NA, 20x LD plan NeoFluar 0.4NA, 50x Epiplan 0.55NA.

### Extra-long working distance air objectives: a mandatory constrain

In high-pressure freezing (HPF) experiments, the primary challenge is to minimize the heat extraction time at a given pressure, in relation to the sample thickness, to facilitate water glass transition faster than ice nucleation. To achieve this, the sample must be thinned as much as possible, with an upper limit of 200 µm^3,5,30–32^. Any addition of material (solid or liquid) increases the heat extraction time, deviating from the fine balance achieved in HPF. To illustrate this delicate equilibrium, it is important to note that vitrification at 2000 bars can only be achieved below -100°C and with a heat extraction rate above 2000K.s^-1^ to transition from the solid to liquid phase (STL) limit around -20°C to below the TTL limit^33^, around -100°C at this pressure ^24,34^. To this end, liquid nitrogen is used, but at 2000 bars, its temperature is no longer -196°C, but only -138°C at best, significantly reducing the temperature gradient. Considering that water turns back to cubic ice around -135°C, at atmospheric pressure, this provides a narrow window in time, pressure, and temperature to achieve vitrification. The presence of immersion media could potentially disrupt this delicate balance.

In designing this project, our main objective was to achieve the fastest time correlation possible. Imaging samples at the highest resolution possible necessitates the use of immersion media. However, removing oil by blotting is slow and rarely complete, with the remaining oil being expelled through the exhaust pipes of the HPM, risking long-term damage and early degradation of an expensive apparatus. Blotting water could result in residual surface water in the micron range, potentially impeding sample cooling. Therefore, we disregarded the option of blotting immersion media early in the project.

Instead, we opted for air objectives and considered the simplest assembly possible using extra-long working-distance objectives. This extended distance provides more space to design the HPF vessel to protect the CryoCapsule, namely the HPF Clamp. The distance also safeguards the objective lens from physical contact with the HPF clamp – which must quickly retract upon HPF trigger – to prevent damage to the objective by scratching the surface (Figure 2C).

The objectives retained on our setup are 5x Fluar 0.25NA, 10x EC plan NeoFluar 0.3NA, 20x LD plan NeoFluar 0.4NA, 50x Epiplan neofluar 0.55NA. Their free working distances are 12.5mm, 5.5mm, 7.9 and 9.1mm respectively.

### Sample vibrations and sharp imaging at 50x: creation of a dedicated high-resolution stage

Operating the system in the aforementioned configuration enables an unprecedented time correlation between live imaging and cryo-immobilization. The mechanical bridge between the high-pressure freezing machine (HPM) and the air-lifted microscope breadboard physically synchronizes the vibrations from both equipment, resulting in sharp images at low magnification (5x) and up to 20x, using a widefield fluorescence microscope, in time and 3D (Z) (Figure 3, Supplementary Figure 1-5).

**Figure 3:**
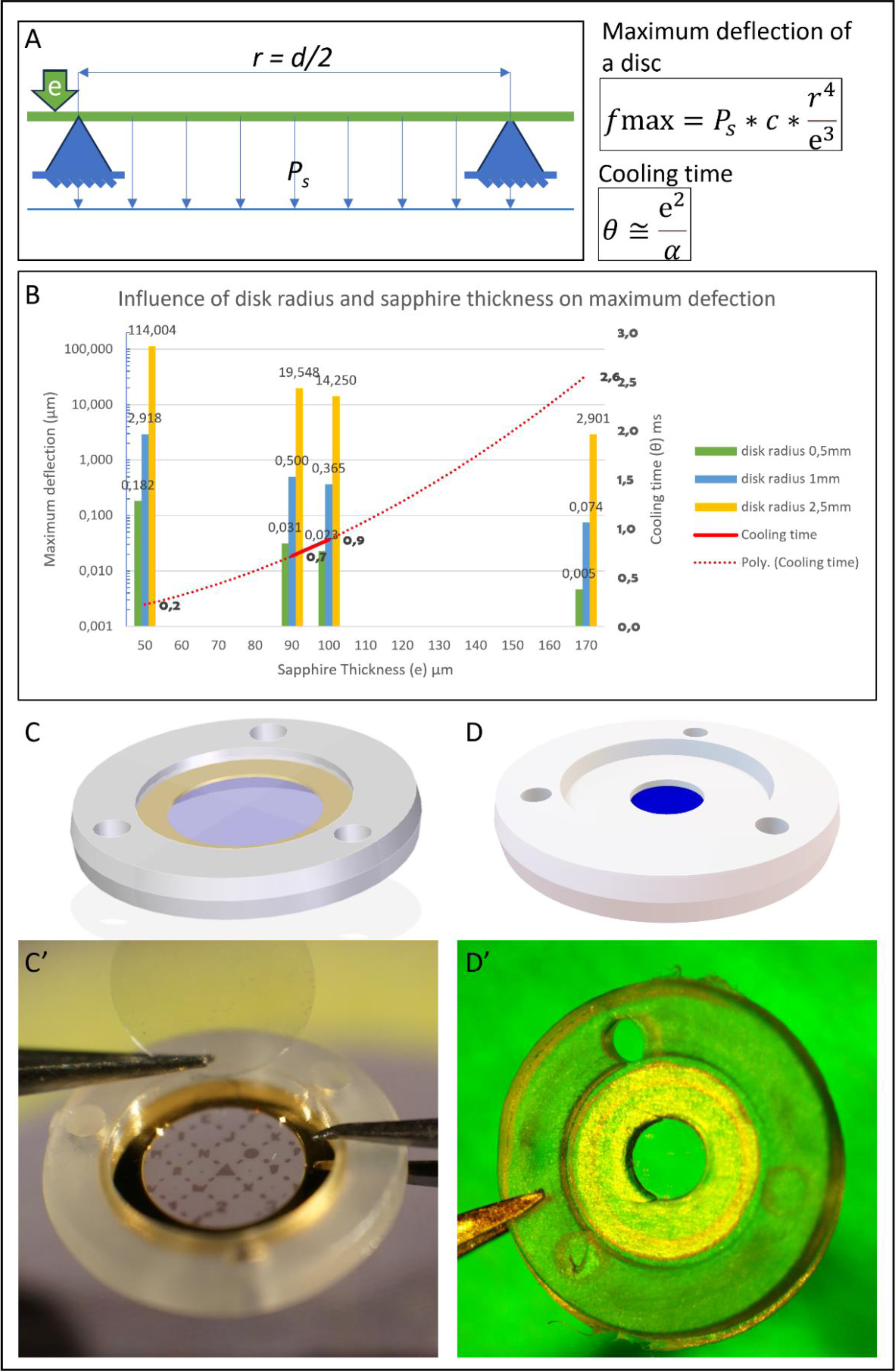
CryoCapsule geometry optimization. A-Schematic representation of the sapphire disc supported by the spacer ring (blue pyramids) and supporting a differential pressure (Ps). The thickness of the sapphire is noted ‘e’, the radius ‘r’ (1/2 diameter ‘d’). Equation 1: simplified equation of the maximum deflextion of a disc supported by a ring. Equation 2: simplified equation of the cooling time of a plate. B-Graphical representation of the two equations afore mentioned. The maximum deflection is represented relative to the ring radius for various standard sapphire thickness. For each thickness, the cooling time is calculated. C-schematic view of the first CryoCapsule. C’-Picture of the first CryoCapsule. The sapphire was 50 µm thick, and the ring was 2 mm in diameter, causing a deflection of nearly 3 µm. D-Schematic view of the CryoCapsule 101. D’-Picture of the CryoCapsule 101. The ring diameter is reduced to 1 mm and the thickness of the sapphire increased to 100 µm, resulting in a deflection of only 23 nm.

However, using a higher magnification objective (50x), obtained images were suboptimal compared to the standalone microscope, where the sample is held on the usual microscope stage. The remaining vibrations between both apparatuses could no longer be compensated by the mechanical bridge, hindering the accurate monitoring of endosome movements using our epifluorescence microscope (data not shown).

To address this issue, we designed a dedicated stage that holds the sample holder containing the CryoCapsule at the focal plane of the microscope, in place of the regular microscope X-Y stage and aligned to the HPM entry site. In this configuration, the imaging stability is comparable to regular imaging conditions (Figure 2, G-K).

To push the sample into the HPM live µ and hook it to the robotic arm for automated vitrification, we added a manual linear transfer arm. This manual operation takes approximately half a second and must be added to the internal transfer of 1.26 seconds mentioned earlier. Therefore, it is reasonable to estimate that the time delay between the last image and vitrification occurs within less than 2 seconds, allowing our technology to address a large span of scientific questions by correlative light and electron microscopy (CLEM). (Video 3)

### Culturing samples in physiological conditions on a transparent but heat conductive and pressure resistant vessel: the CryoCapsule ^6,7,12–14^

HPF aims at vitrifying specimens within a thickness range of 5 µm to 200 µm, surpassing the capabilities of conventional plunge freezing. Traditionally, samples are placed between two metallic carriers, typically made of aluminium or copper. These carriers facilitate the transmission of pressure without exerting uneven compression on the specimen. Additionally, they dissipate heat to achieve vitrification. However, due to the opaque nature of metal, the pre-loaded sample within an HPF apparatus is unobservable.

In light of this limitation, the implementation of sapphire emerged as a compelling solution^35^. Sapphire (aluminium oxide) has remarkable properties including high thermal conductivity, mechanical robustness, and excellent optical properties ^6,35–37^. Thus, it stands as the optimal material for fabricating the sample vehicle in HPF. Building upon the foundational design of the CryoCapsule^25,26^, we have taken evolutionary strides to enhance its features. In contrast to the original design which employed a 50 µm thin sapphire – a choice supported by initial publications advocating its utilization ^6,35,36^– we found this thickness to exhibit excessive flexibility (Figure 3, equation 1. maximum flexion point 2.9 µm, nearly a third of the cell’s thickness). This leads to uneven deformations during the HPF process and subsequent detachment of cells from the sapphire disc. Conversely, at a thickness of 160 µm, rigidity ceased to be problematic (74 nm). However, the freezing reliability became inconsistent in our setup (data not shown). This inconsistency may be attributed to reduced cooling efficiency. The cooling time increases by the square of the thickness “e”, (Figure 3, equation 2) (from 0.22 ms to 2,56 ms). A 100 µm thick sapphire disc has a cooling time of 0.89 ms.

The diagram (Figure 3, B) illustrates the influence of the disk radius and the sapphire thickness in the maximum deflection and presents the cooling time according to the sapphire thickness.

To further refine the CryoCapsule design, several key modifications were made. The cavity diameter was reduced from 2 mm to 1 mm, improving two aspects: 1. a narrower sample area is easier to correlate in the electron microscope, minoring the need for carbon landmarks. 2. it further reduces the 100 µm thick sapphire disc maximum flexion down to 23 nm. Simultaneously, the capsule’s depth was increased from 50 µm to 100 µm, thereby broadening its applicability to accommodate various specimens such as biopsies, organoids, and small model organisms (Figure 3, C-D’).

### Freeze Substitution and embedding

Following vitrification, the samples underwent freeze substitution using an FS-8500 RMC (Boeckeler Instruments, Tucson, Arizona) in a solution mix detailed in Table 1. The freeze substitution process followed a temperature ramp program as outlined below:

For the standard freeze substitution protocol:

-18 hours at -90 degrees Celsius
-15 hours from -90 degrees to -60 degrees Celsius (with a temperature increase of 2°C per hour)
-8 hours at -60 degrees Celsius
-15 hours from -60 degrees to -30 degrees Celsius (with a temperature increase of 2°C per hour)

Upon completion of the program, the samples were kept on ice for 1 hour, followed by three washes in acetone. Subsequently, the samples were embedded in EPON 812 resin (EmBed-812 Embedding Kit, Electron Microscopy Sciences, Hatfield, Pennsylvania, USA) through a series of steps below.

Preparation of EPON 812: Mix1 (2 mL Embed-812, 3.1 mL Dodecenyl Succinic Anhydride) and Mix2 (2 mL Embed-812, 1.7 mL Methyl-5-Norbornene-2, 3-Dicarboxylic Anhydride) were mixed just before the use and added 170 µL DMP-30. We used this solution as 100% resin.

Dehydration and embedding step: 2 hours each at 30%, 60%, and 100% resin concentration in acetone. At 100% was repeated twice. The embedding process concluded with 48 hours of polymerization at 45°C. 60°C is recommended for normal polymerisation. In our case, an oven fault limited the temperature to 45°C. For this reason, we extended the polymerisation time, after which no impact of the lower temperature on sample preservation was observed.

### Ultra microtome

70 nm to 80 nm thin sections were made on an Ultracut S (Leica) with an ultra 45° then collected on homemade formvar coated mesh grids, or formvar coated slot grids.

### Transmission Electron Microscope Imaging

TEM imaging was performed with a Tecnai 12 transmission electron microscope, 120 kV (FEI) equipped with a CCD camera (OneView 4Kx4K Gatan) controlled by GMS software.

### Individual sample datasheet

All presented material is individually registered into F°Low ^38^, our project management software, and their unique sample preparation protocol is synthesized into a 2-page PDF for easy reproduction of the protocol presented in this article. All twenty files are presented as supplementary material (F0Low-datasheets.zip).

### Workflow

#### In practical implementation, the initial step is loading the sample into the CryoCapsule

*Note: The CryoCapsule is delivered with a bare sapphire disc in it, without any coating. It is left to the experimenter to do their usual coating, appropriate to grow their cell line of interest. In this study, the cell line used required no prior coating. Adhesion is not enhanced by poly-L-lysin or fibronectin as they perturb the normal behaviour of the cell*.

For the culture of cell monolayers, the CryoCapsules are put within a larger Petri dish wherein cells are cultured^25,26^. After seeding, the cell adhesion and growth are meticulously monitored within a controlled culture environment in each CryoCapsule. On the day of the experiment, the Petri dish containing one to three CryoCapsules is transported to the HPF room. Here, the samples are examined using the standalone microscope configuration (Figure 2A, video 1).

In this setup, the microscope serves as the frame of reference. The manual microscope stage is employed to navigate across the CryoCapsule (video 1: 0’17). Once the cell of interest reaches the optimal stage for vitrification, a seamless transition is performed. The CryoCapsule is transferred to the HPM clamp, situated at the robotic arm’s tip (video 1: 1’13). Concurrently, the microscope is transformed to fit to the HPM Live µ (Figure 2B, video 1: 0’40, 1’30).

This entire process takes less than a minute, if executed by two experimenters simultaneously (experienced user can do it alone in one minute, video 1: 0’40 to 1’40). Subsequently, the cell of interest gets positioned using a 5x magnification, followed by precise centring utilizing the X-Y shifting table beneath the microscope (Video 1, 1’00”) as the HPM now serves as the frame of reference. A suitable magnification, usually up to a 20x air objective, is chosen, enabling real-time observation up to 10 minutes. Throughout this observation, the HPM remains poised to initiate vitrification whenever needed.

Upon the detection of the event of interest, the experimentalist can trigger the vitrification process. This can be accomplished by either pressing a foot pedal or using the HPM screen interface. The cryo-immobilization process is achieved 1.26 seconds after the initiation command, ensuring swift and precise capture of the biological event (Video 1: 3’07).

In the case where a higher magnification is required, such as 50x for endosome tracking, we used a microscope stage designed for this aim (Figure 2, G; Video 2). We push the sample into the HPM at the end of the image acquisition (Figure 2, J; Video 2: 0’56’’77). The sample holder is secured to the robotic arm, and simultaneously, the HPF cycle is triggered (Figure 2, K; Video 2: 0’57’’42). This cycle takes less than 2 seconds altogether.

We completed the experiments by freeze-substituting the samples for later electron microscopy analysis and CLEM registration.

## Results

### Reliability of the workflow

One major challenge in live-HPF workflow is to limit sample damage or displacement during vitrification while preserving physiological conditions. The shock of liquid nitrogen reaching the sample at a pressure rise rate above 100 bars per millisecond can cause support distortion within its flexibility range. This is the main source of the sample damage and displacement, and it needs to be reduced. To address this challenge, we optimized our technological tandem, CryoCapsule/HPM Live µ, to achieve the highest success rate possible, in absence of adherence treatment.

We cultured cells stably expressing tau-GFP proteins, labelling microtubule. Various confluences were tested to account for individual adherence and the risk of large group detachment. In these experiments, we cultured cells on CryoCapsules and mapped them at low magnification in the petri dish prior to loading into the CryoCapsule clamp. After confirming that the cells were well distributed, each CryoCapsule was closed by an extra sapphire disc, loaded in the HPF clamp, and inserted at the tip of the robotic arm. Fluorescence images were acquired at 5x, 10x, and 20x magnification.

We note that the images taken before loading were sub-optimal, as the light path went through the plastic petri dish, the cell culture medium, and the support sapphire disc of the CryoCapsule, but this does not prevent cell mapping and ROI identification (Figure 4A-D). However, once loaded in the HPF clamp, the light path of the air-objective went solely through the support sapphire disc of the CryoCapsule, resulting in a better signal-to-noise ratio and sharper identification of the microtubule network (Figure 4, E, F, I, M, Q). Importantly, no cells were lost or detached from the CryoCapsule during the process. It gave us confidence in sample tracking through this multiple-step workflow (Figure 4, B, E).

**Figure 4:**
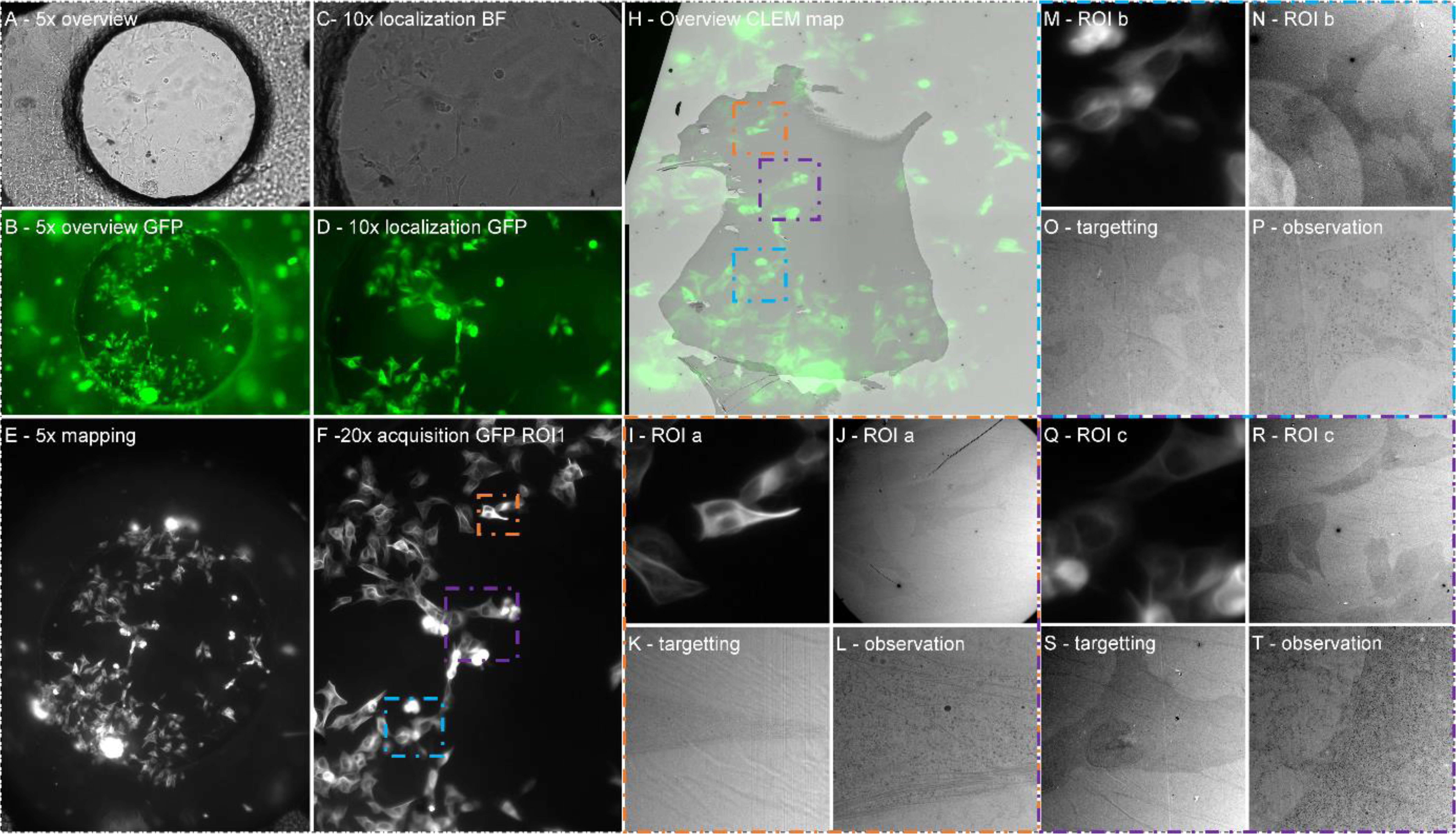
Full Live-CLEM tracking of multiple cells. A-Observation of the CryoCapsule content at low magnification on the microscope in Standalone configuration. B-Overview of the SY5Y-SH cells expressing Tau-GFP. C-Localization of a group of cells of interest. D-Observation of the group of cells of interest in bright field. E-Overview of the CryoCapsule with the integrated microscope. F-live imaging of cells of interest at 20x, ROI in orange (a), blue (b) and purple (c) are highlighted in H. H-Overlay of the first section by TEM with the 5x mapping. ROI in orange (a), blue (b) and purple (c) are magnified in I-T. I-L-ROI a (Orange), Cell of interest (I) is localised (J), targeted (K) and the ROI is observed (L). All 3 ROIs (a, b, c) are presented in a similar way to illustrate our ability to retrieve all cells in one CryoCapsule.

The CryoCapsule was vitrified, freeze-substituted, and embedded in epoxy resin. The first sections of each block were collected, trying to be as parallel as possible to the block face to rapidly obtain a full section of the block. As the CryoCapsule has a diameter of 1 mm, it is possible to section the entire block face on an ultramicrotome and collect the entire section on a slot grid. This strategy allows for matching each cell to its live fluorescent detection.

We first generated an overview CLEM map to find the cells we identified by live imaging (Figure 4, H) and then navigated to various areas to observe microtubule bundles at the plasma membrane (Figure 4, J, N, R). It is important to note that across several experiments, the cells retrieval occurs in an ‘all or nothing’ fashion. If the CryoCapsule has encountered a damage during sample preparation, then all the cells are lost or moved, across the entire surface of the CryoCapsule. If the integrity of the CryoCapsule is not compromised during the workflow, then all the cells are retrieved in the electron microscope, at the expected position. Even loosely attached mitotic cells were retrieved at their expected position (ref: R221 article). To demonstrate the reliability of the workflow, the 6 CryoCapsule that were vitrified on that experiment are presented (Figure 4, Supplementary Figure 1-5), resulting in 100% retrieval in that experiment. Figure 5 further demonstrate the reliability of our workflow, and two extra independent experiments are presented in the manuscript about the R221 resin. Systematic displacement of the cells was screened by bloc-face in-resin fluorescence microscopy over two years, confirming the reliability of the workflow (data not shown) opening the path to more advanced sample preparation.

**Figure 5:**
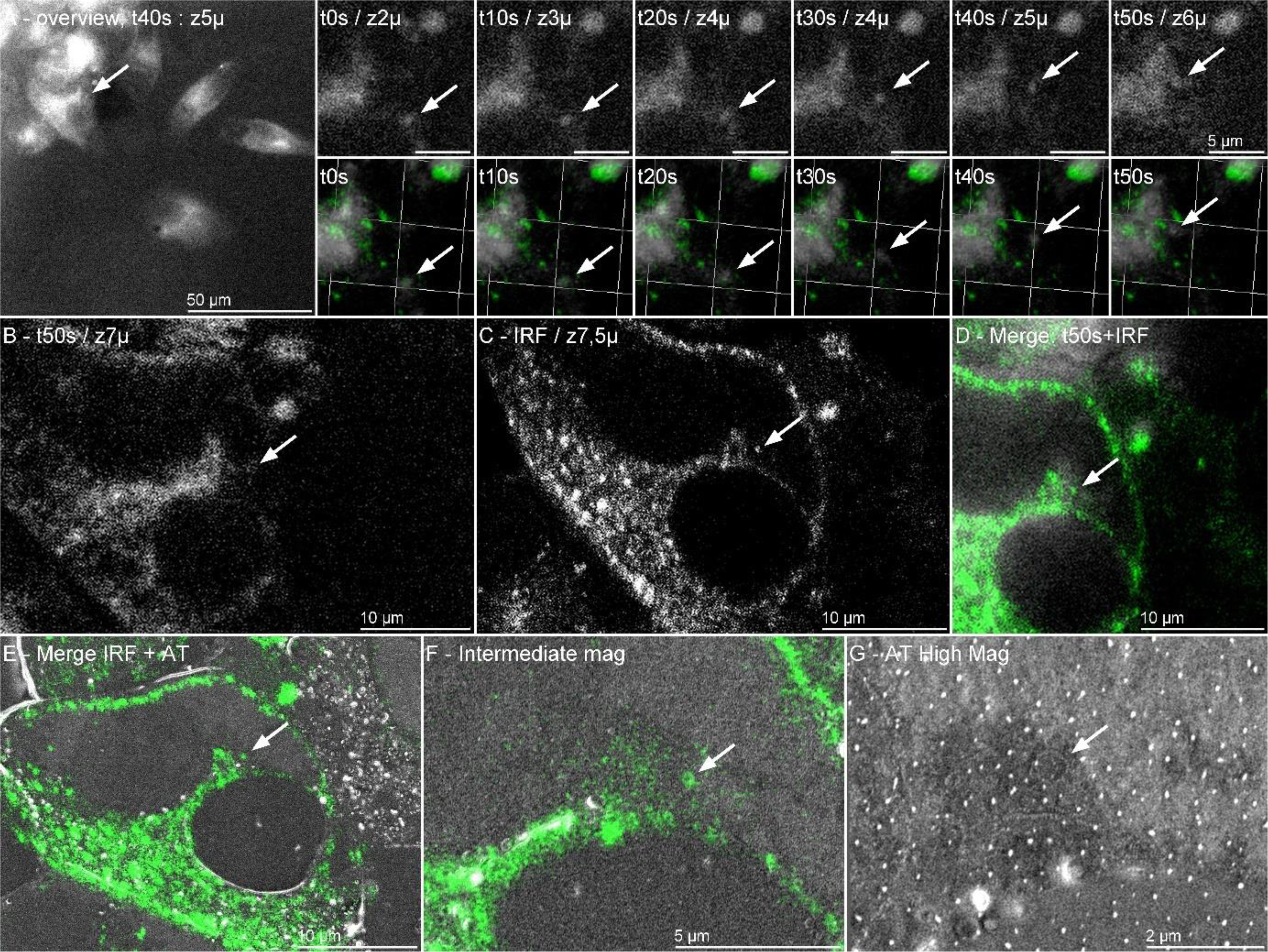
Full Live-CLEM tracking of a dynamic ensodome. A-at 50x, several cells are visible in the field of view. One has an isolated endosome that is traceable during the full imaging procedure. We see that it navigates approximately 5 micrometres in X, Y and across 4 micrometres in Z during the imaging period of 60 seconds at 10 s interval. Registering the live data with the confocal stack, we observe that the fluorescent signal moves towards the isolated fluorescent signal used as landmark to search for the endosome in the array tomography stack. The arrow points towards the endosome. B-Last image before vitrification. C-Isolated slice from the confocal stack. D-The last frame of the live dataset is registered to the confocal stack to identify the isolated endosome, tracked live. E-An Array Tomography 3D stack is registered to the confocal stack. F-at intermediate magnification, we localize the endosome. G-At higher magnification, the endosome is retrieved at the expected X-Y coordinates, but 5 slices further in Z, illustrating a registration discrepancy in Z of 500 nm.

### Application to a dynamic biological structure: high spatial and temporal resolution of a Full CLEM Workflow

The endosomal network regulating intracellular transport is highly developed and intricated in melanocytic cells, where effective melanosome transport is needed. Dynamic, it moves in the range of hundreds of milliseconds, appear as rounded structures with tubulation, and are easily identified in the electron microscope as they are relatively large but discrete structures (150 to 500 nm diameter). Tracking them live before fast immobilization and retrieving them at the electron microscope level is essential to decipher their function^39^. However, correlating one individual compartment from the live epifluorescence microscopy dataset with serial section volume transmission electron microscopy images is challenging. Serial thin section accurate collection associated to the risk of loss and distortions leads to a very low throughput. Serial section electron tomography is also laborious. In the past, we addressed this challenge by on-section fluorescence microscopy to narrow structural assignments ^39–41^ as volume electron microscopy was not as widely spread as today.

We cultured melanocytic cells (MNT1) stably expressing an endosomal marker, syntaxin-13-GFP, at endogenous levels (MNT1-Stx13-GFP), in the CryoCapsule and imaged them using the high-resolution stage. This secondary equipment allowed us to obtain images without blurring caused by asynchronous vibrations. We tracked individual endosome for one minute, every 10 seconds, at 50x across 15 µm (Z-stack 15 x 1 µm) from isolated cells prior to vitrification.

3D resolution in epifluorescence is diffraction limited. This is further compounded using long working distance objective lenses imposed by the system. However, it remains sensitive enough to detect an event in the X-Y plane, as epifluorescence also induces a deeper focal depth, requiring fewer Z-stacks to localize and track an event of interest.

We identified a single endosome travelling around the nucleus. The 3D+time dataset allowed us to track the endosome in the cell volume (Figure 5A, T0 to t50s). The endosome navigates across approximately 5 µm in X and Y and across 4 µm in depth (from Z2 to Z6) over a duration of 60 seconds (10 seconds intervals). At the end of the acquisition, we rapidly initiated the vitrification (in about 2 seconds). In this experiment, the freeze-substitution cocktail contained only traces of uranyl acetate (0.1%) and water (5%) to preserve the faint fluorescence. Following embedding in R221, we imaged the sample bloc through a glass bottom petri dish, at high resolution (63x, oil immersion) using a laser scanning confocal microscope (SP8, Leica, Germany). This fixed confocal dataset was registered to the live dataset to relocate the endosome of interest. The precise re-localization in X and Y of the endosome, isolated from other fluorescent structures, confirms the fast cryo-immobilization after live imaging. We re-estimated the endosome depth to be 6 µm deep in the cell according to the confocal. 105 Serial sections of a 100 nm were collected on two wafers and first imaged at low magnification to reconstruct a volume of the cell ^42^ that we registered to the confocal stack. the sections above and below 6 µm depth were imaged to locate the endosome of interest (approximately section 45 to 70). 30 sections were imaged at higher magnification in the region of interest to reconstruct a volume of interest, that was again registered to the confocal stack The precision of localization is accurate in X and Y, but we found the endosome on section 53 instead of the expected section 60.

Altogether, this demonstrates the comprehensiveness of the workflow we established, encompassing sample culture, sample live imaging, vitrification and post-fixation localization at high resolution to ultimately image by electron microscopy of the unique structure of interest.

## Discussion

The concept of live cell imaging rapidly followed by HPF for CLEM strategies is not novel and was first proposed by Paul Verkade in 2008 with an innovative sample loading strategy ^1^. The supporting sapphire disc was held into a perforated membrane carrier to comply with the EMPACT-2 equipped with the rapid transfer system (RTS, Leica microsystem). Despite the broad enthusiasm of the electron microscopy field for this promising technology, very few laboratories were able to establish this method as a routine. The following High Pressure Freezing machines on the market (HPM100, EM ICE, Leica Microsystem; Wohlwend compact 03, Wohlwend) were also holding this strategy as a promise, but to our knowledge, no publication demonstrates the reproducibility of the technique on any of the afore mentioned apparatus. The technological challenges associated with live-HPF CLEM include *live imaging quality, reproducibility of time transfer, sample preservation, vitrification quality, and cells retrieval reliability*. Our study addresses these challenges by developing an automated and standardized workflow.

With the first version of the CryoCapsule^25,26^, we already addressed some of the challenges: live imaging directly across the bottom sapphire disc, without any immersion, offers sufficient detection capabilities to localize and monitor an event of interest, accepting some compromise on the resolution in comparison to state of the art live imaging (oil immersion objectives, confocal or super-resolution imaging). We also demonstrated the vitrification quality, and to some extent, the cells retrieval was addressed but required some further improvements.

### Time transfer monitoring could only be addressed by automating this step and therefore combining the microscope and the high-pressure freezer

Last but not least, is it notorious that sapphire disc freezing is a delicate process, and many vitrification cycle led to sapphire rupture, with associated sample loss and equipment damages (sapphire shrapnel dispersion in the exhaust pipes).

We therefore conceived our own High-Pressure Freezer, in partnership with ABRA Fluid AG, the historical HPM010 manufacturer, to get a full control on the design and evolution of the various elements composing this complex workflow^24^.

### Sample transfer automation

Our primary goal was the automation of the sample transfer with the pre-existing CryoCapsule. From the live imaging position, the sample must be retracted and inserted into the HPF chamber in the fastest way possible. We opted for a single translation rotation move, as short as possible to minimize time transfer and achieved 1.27 second from the trigger to the vitrification ^24^. Study of phenomenon faster than that would require either protocol adaptation or redesigning the system.

### Light microscope design

To prototype our workflow assembly, we used an epifluorescent Zeiss Axiovert 200M. On intensely but finely labelled structures such as the microtubule network, we can identify isolated microtubule bundles. Furthermore, it is sensitive enough to capture individual endosome movement, labelled with Syntaxin-13-GFP expressed at endogenous level, thanks to the high-resolution stage and despite the use of a long working distance 50x air objective with a numerical aperture of 0.5. In practice, this proves our set-up to be sufficient to demonstrate our ability to track dynamic individual cellular event. Obviously, the use of a more advanced microscope set-up such as a spinning disc confocal microscope, adaptive optics or an Airyscan microscope (LSM900 Airyscan, Zeiss, Germany) will benefit to the live imaging detection, especially compensate for spatial resolution. The use of advanced illumination technology also opens the way for photo manipulation (optical tweezers, laser dissection, photo activation, photobleaching), broadening significantly the application spectrum of live-HPF CLEM.

### Cost optimization

High Pressure Freezing is the entry point of long and tedious workflows, and never appears to be the bottle neck of any electron microscopy pipeline. Therefore, associating the HPM Live µ to a high-end microscope may appear difficult to justify to funding organizations. This is why we set the microscope on rails to allow independent use of the LSM900 in imaging facilities. This rail system was developed specifically for the integration with the HPM Live µ, including a specifically manufactured vibration free table.

### Experiment reliability

To improve the sample preservation, many aspects were considered. Two major issues are frequently reported when discussing about live-HPF CLEM: sapphire rupture and cells displacement. Both events are dramatic as they cause apparatus damage and sample loss, often hindering weeks of cell biology sample preparation. The liquid nitrogen flow at such high pressure (2000 bars) and speed (200 bars.ms^-1^) is turbulent, causing uneven pressure on both sides of the sapphire assembly. Flexion occurs up to cells detachment or sapphire rupture. To account for this, scientists have created sapphire assembly including at least one aluminium support^2^. This system is more resistant and prevents rupture but is unable to completely remove the effect of the flexion. As a consequence, cells displacement is often observed, and scientist compensate by mapping the entire sapphire substrate and acquire multiple cells in locations apart to get a chance to save one sample Another attempt is coating the sapphire disc with poly-L-lysine that glues the cell to the surface, but it alters the behaviour of the cells in comparison to regular light microscopy analysis. Furthermore, the manual assembly of the sapphire is complicated and often expands sample loading time or potential exposure to air. To facilitate sapphire manipulation, manufacturers proposed increasing its diameter and consequently increased its thickness. This resulted in impairing vitrification quality. The strategy with the CryoCapsule takes all these reversely. We calculated the optimal dimensions of the sapphire substrate relative to its thermal and mechanical properties. This led to reducing the sapphire disc diameter, closer to the initial work from Verkade et al. To ease the sapphire disc manipulation, the sapphire is embodied in a plastic frame. The natural elasticity of the plastic frame damps the turbulences during the HPF shot, contributing simultaneously to avoid sapphire deformation and cells detachment. The necessary increase in thickness from the original CryoCapsule, to account for the deformation (varying by the cube of the thickness, eq. 1), is defined relative to the maximum heat transfer (varying by the square of the thickness eq. 2). The crossing of both curves is near 85 µm. We opted for a 100 µm thick sapphire, that keeps the flexion below 25 nm, the diameter of a microtubule.

This combination of technical optimization allows us to retrieve 100% of the cells (unless a damage occurred in the sample preparation) with a very high quality of vitrification across the whole surface of this newly designed CryoCapsule.

### 3D correlation

The final step of this comprehensive workflow is the location of the specimen tracked live into the electron microscope. The inherent limited resolution of our epifluorescence microscope impairs the capacity to locate a fine structure in Z. This imposes a systematic use of volume electron microscopy techniques to ensure retrieving the isolated element. The delay and costs to access these equipment’s are not always compatible with the frame of a project. In-resin fluorescence would compensate for the low resolution of the microscope and generate a high-resolution map of the processed sample (already dehydrated, stained and embedded). Imaging this same block by electron microscopy would improve image registration. Here, R221 plays a central role as it simultaneously allows in-resin fluorescence strategies and volume EM imaging techniques.

This full workflow, from live imaging to volume EM is made possible by the combination of three central technologies: the CryoCapsule, the HPM Live µ and the R221, filling the sample preparation gaps between all imaging steps.

## Supplementary Data

Video 1: https://youtu.be/6QnGl6kPegs

Video 2: https://youtube.com/shorts/qkACmvkdOLQ

## Supplementary Figures

**Sup Figure 6:**
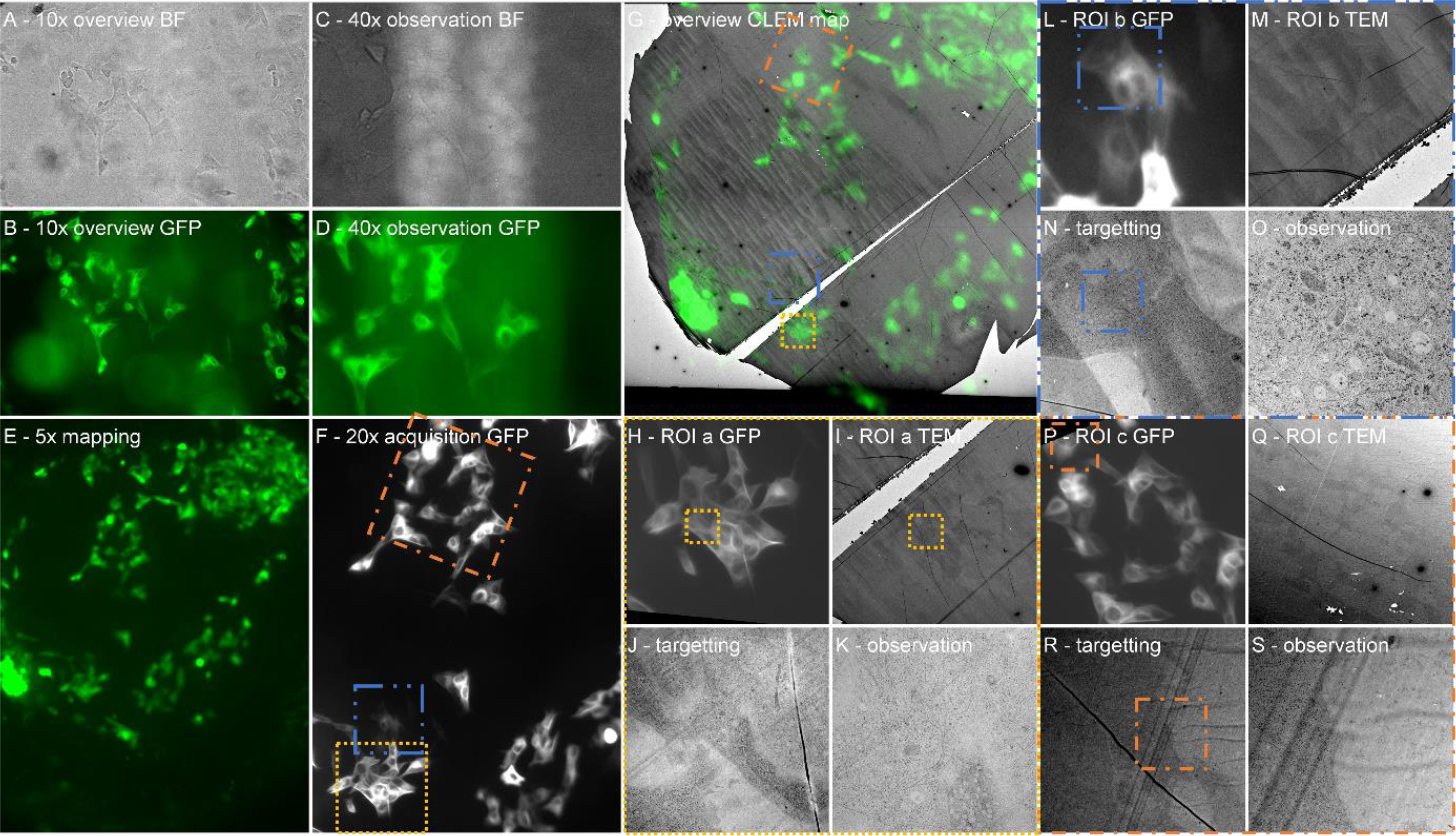
Full Live-CLEM tracking of multiple cells – Capsule N°2. A-Observation of the CryoCapsule content at low magnification on the microscope in Standalone configuration. B-Overview of the SY5Y-SH cells expressing Tau-GFP. C-Localization of a group of cells of interest. D-Observation of the group of cells of interest in bright field. E-Overview of the CryoCapsule with the integrated microscope. F-live imaging of cells of interest at 20x, ROI in orange (a), blue (b) and purple (c) are highlighted in G. G-Overlay of the first section by TEM with the 5x mapping. ROI in orange (a), blue (b) and purple (c) are magnified in H-S. H-K-ROI a (Orange), Cell of interest (I) is localised (J), targeted (K) and the ROI is observed (L). All 3 ROIs are presented in a similar way. This illustrates our ability to retrieve all cells in every CryoCapsule.

**Figure 7.**
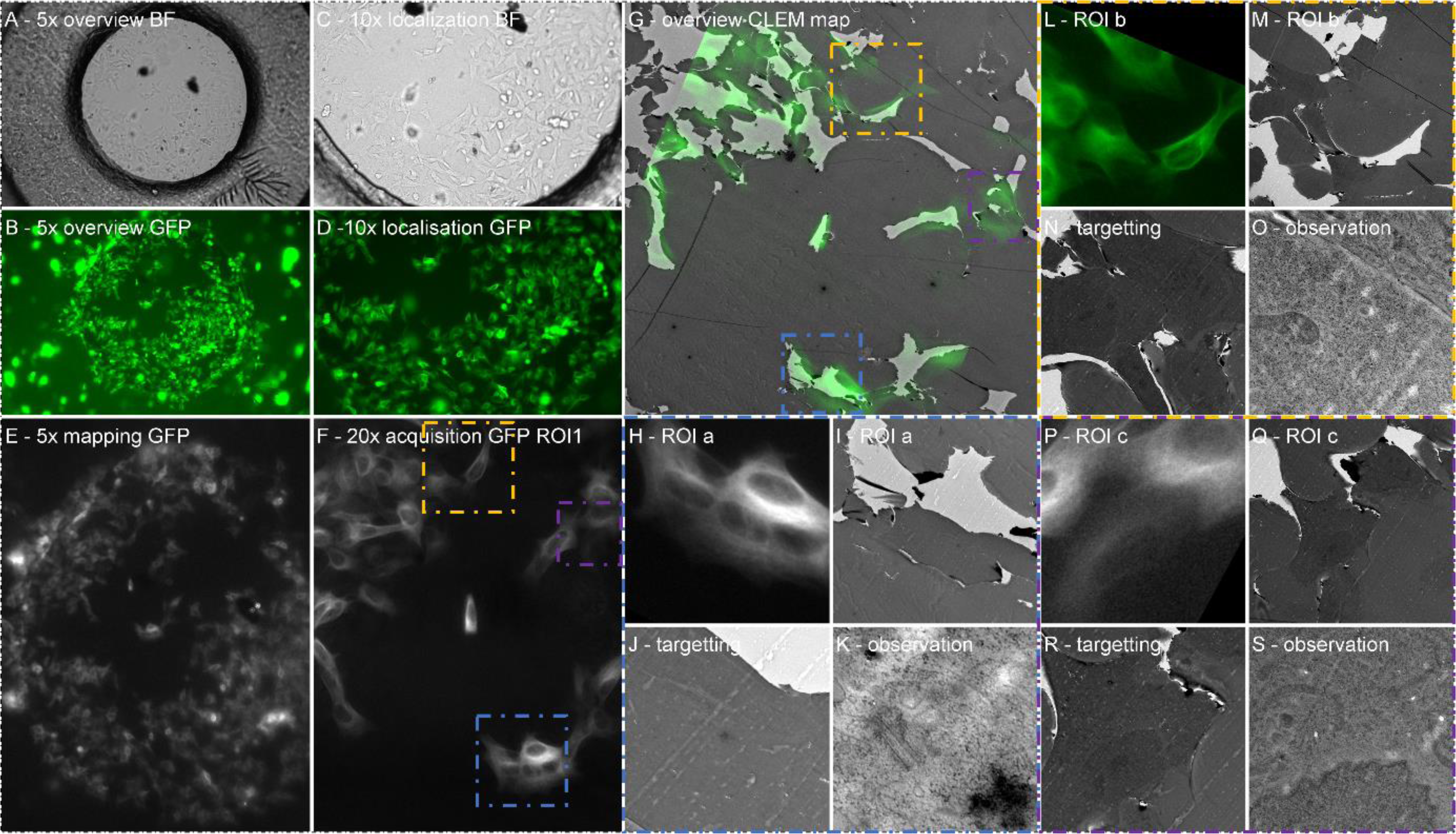
Full Live-CLEM tracking of multiple cells – Capsule N°3. A-Observation of the CryoCapsule content at low magnification on the microscope in Standalone configuration. B-Overview of the SY5Y-SH cells expressing Tau-GFP. C-Localization of a group of cells of interest. D-Observation of the group of cells of interest in bright field. E-Overview of the CryoCapsule with the integrated microscope. F-live imaging of cells of interest at 20x, ROI in orange (a), blue (b) and purple (c) are highlighted in G. G-Overlay of the first section by TEM with the 5x mapping. ROI in orange (a), blue (b) and purple (c) are magnified in H-S. H-K-ROI a (Orange), Cell of interest (I) is localised (J), targeted (K) and the ROI is observed (L). All 3 ROIs are presented in a similar way. This illustrates our ability to retrieve all cells in every CryoCapsule.

**Figure 8.**
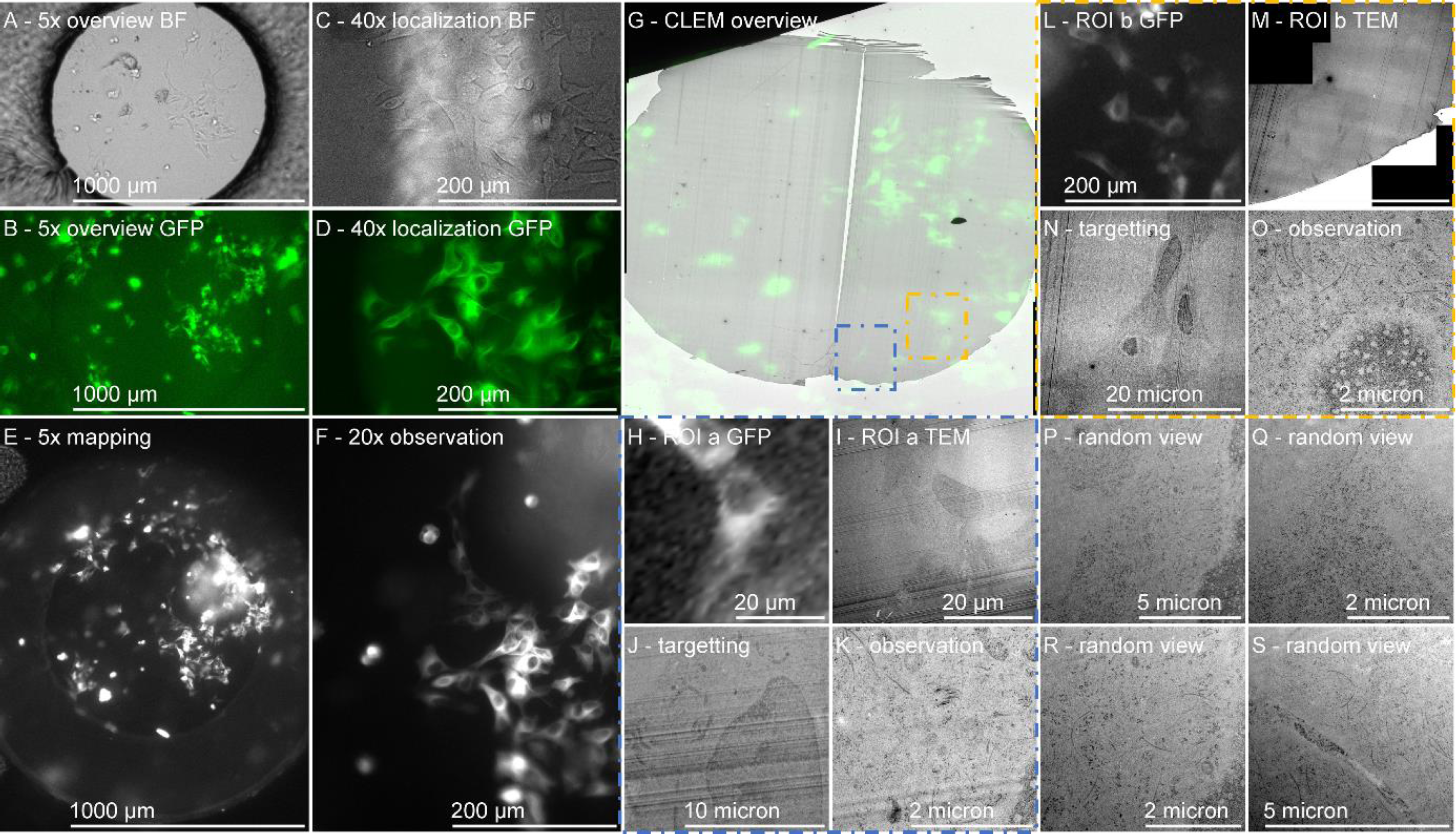
Full Live-CLEM tracking of multiple cells – Capsule N°4. A-Observation of the CryoCapsule content at low magnification on the microscope in Standalone configuration. B-Overview of the SY5Y-SH cells expressing Tau-GFP. C-Localization of a group of cells of interest. D-Observation of the group of cells of interest in bright field. E-Overview of the CryoCapsule with the integrated microscope. F-live imaging of cells of interest at 20x, ROI in orange (a), blue (b) and purple (c) are highlighted in G. G-Overlay of the first section by TEM with the 5x mapping. ROI in orange (a), blue (b) and purple (c) are magnified in H-S. H-K-ROI a (Orange), Cell of interest (I) is localised (J), targeted (K) and the ROI is observed (L). All 3 ROIs are presented in a similar way. This illustrates our ability to retrieve all cells in every CryoCapsule.

**Figure 9.**
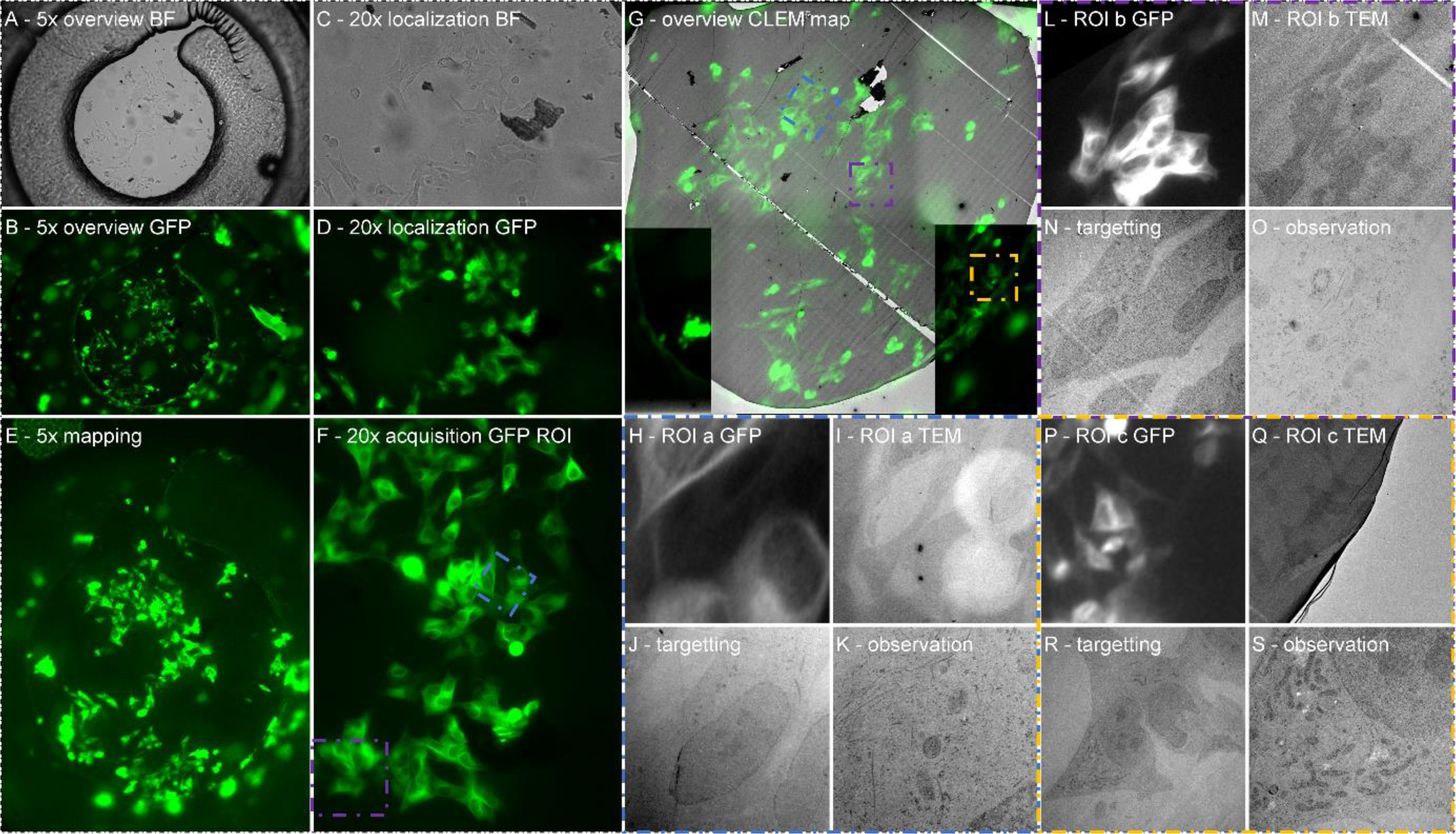
Full Live-CLEM tracking of multiple cells – Capsule N°5. A-Observation of the CryoCapsule content at low magnification on the microscope in Standalone configuration. B-Overview of the SY5Y-SH cells expressing Tau-GFP. C-Localization of a group of cells of interest. D-Observation of the group of cells of interest in bright field. E-Overview of the CryoCapsule with the integrated microscope. F-live imaging of cells of interest at 20x, ROI in orange (a), blue (b) and purple (c) are highlighted in G. G-Overlay of the first section by TEM with the 5x mapping. ROI in orange (a), blue (b) and purple (c) are magnified in H-S. H-K-ROI a (Orange), Cell of interest (I) is localised (J), targeted (K) and the ROI is observed (L). All 3 ROIs are presented in a similar way. This illustrates our ability to retrieve all cells in every CryoCapsule.

**Figure 10.**
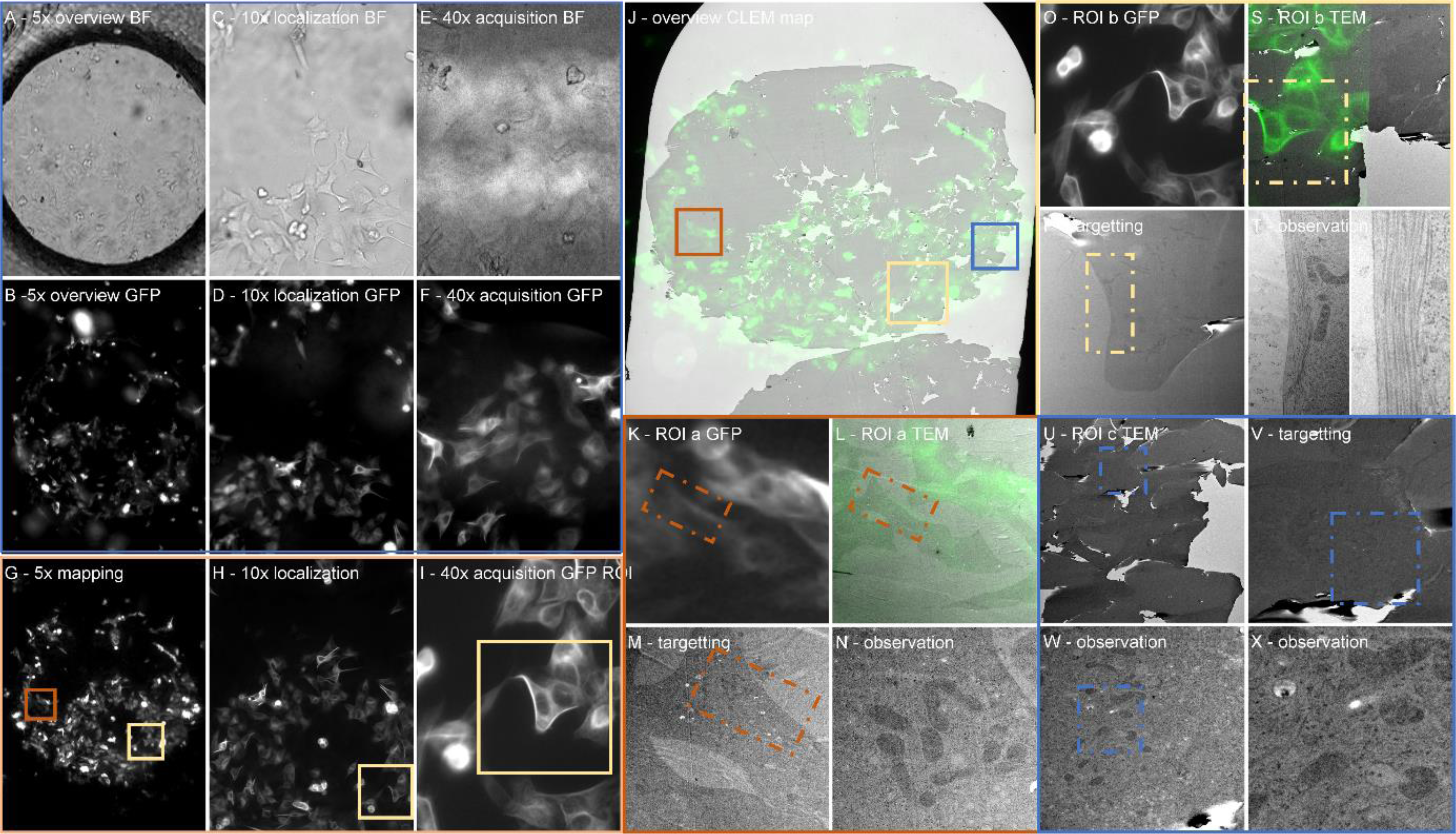
Full Live-CLEM tracking of multiple cells – Capsule N°6. A-Observation of the CryoCapsule content at low magnification on the microscope in Standalone configuration. B-Overview of the SY5Y-SH cells expressing Tau-GFP. C-Localization of a group of cells of interest. D-Observation of the group of cells of interest in bright field. E-Overview of the CryoCapsule with the integrated microscope. F-live imaging of cells of interest at 20x, ROI in orange (a), blue (b) and purple (c) are highlighted in H. H-Overlay of the first section by TEM with the 5x mapping. ROI in orange (a), blue (b) and purple (c) are magnified in I-T. I-L-ROI a (Orange), Cell of interest (I) is localised (J), targeted (K) and the ROI is observed (L). All 3 ROIs are presented in a similar way. This illustrates our ability to retrieve all cells in every CryoCapsule.

